# Chronic pain is linked to a resting-state neural archetype that optimizes learning from punishments

**DOI:** 10.1101/2025.07.11.664303

**Authors:** Francesco Scarlatti, Ludovic Dormegny-Jeanjean, Roman Schefzik, Tobias Banaschewski, Arun L. W. Bokde, Rüdiger Brühl, Sylvane Desrivières, Hugh Garavan, Penny Gowland, Antoine Grigis, Andreas Heinz, Jean-Luc Martinot, Marie-Laure Paillère Martinot, Eric Artiges, Dimitri Papadopoulos Orfanos, Luise Poustka, Michael N. Smolka, Sarah Hohmann, Nathalie Holz, Nilakshi Vaidya, Henrik Walter, Robert Whelan, Gunter Schumann, Frauke Nees, Emanuel Schwarz, Martin Löffler, Jack R. Foucher, Herta Flor, IMAGEN Consortium

## Abstract

Chronic pain is a leading cause of disability, yet its underlying susceptibility traits remain unclear. Disorders like chronic pain may stem from extreme neural types, or archetypes, optimized for specific cognitive strategies and reflected in patterns of resting-state networks. Here, we examined a sample from the general population (*n* = 892) and three clinical samples with subacute back pain (*n* = 76), chronic back pain (*n* = 30), and treatment-resistant depression (*n* = 24). Using the sample from the general population, we found three neural archetypes that prioritize different cognitive strategies. Clinical pain samples, compared to the sample from the general population, mapped close to an archetype optimized for punishment learning (Archetype P). We replicated these results by recomputing the archetypes starting from the clinical pain samples, additionally revealing an association between Archetype P and pain severity. These findings suggest a neural-cognitive trait underlying susceptibility to chronic pain.

## Introduction

Chronic pain is among the primary causes of disability worldwide. Over the nearly three-decade period from 1990 to 2017, two out of the four leading causes of years lived with disability were low back pain and headache disorders (*1*). Instrumental learning, a process in which organisms associate behaviors with their outcomes, plays an important role in the development of chronic pain. This occurs, in part, because individuals develop behaviors to avoid or escape from pain such as reduction of movement or medication use, which may themselves enhance pain via reinforcement learning (*2–7*). Significant efforts have been made to identify the neural and cognitive traits that predispose individuals to chronic pain, including those linked to instrumental learning, such as sensitivity to rewards and punishments (*8–11*). However, although several of these traits have been identified, we still lack a clear understanding of how they are interconnected.

Neural and cognitive traits evolve in response to environmental demands, resulting in brain activity patterns that reflect cognitive strategies adapted to those demands (*12–15*). Also during the resting state, when awake and with their eyes closed, people simulate scenarios and engage in a variety of mental activities, from planning the future to vivid internal conversations, all of which are shaped by personal goals, challenges, or the immediate context. Specifically, lying still in a scanner for functional magnetic resonance imaging (fMRI) can be stressful and people may cope with this situation by adopting different cognitive strategies. These resting-state mental activities are underpinned by resting-state networks (RSNs), systems of interconnected brain regions that function together during rest. For example, if someone frequently engages in planning the future, a greater proportion of their RSN activity may be attributed to the default mode network (DMN), which is associated with such mental processes (*16*).

This relationship between RSN patterns, cognitive strategies, and environmental demands may, provide a foundation for identifying neural-cognitive traits relevant to clinical disorders. These neural-cognitive traits may be best conceptualized as trade-offs between extreme neural types, called archetypes, each optimized for a particular cognitive function. Because real-life demands require balancing multiple cognitive functions, most individuals from the general population likely fall between these extremes, adopting an intermediate form of cognition. However, those with highly specialized cognitive profiles, optimized for a single cognitive strategy at the expense of others, may be more susceptible to certain disorders, including chronic pain (*8*, *17*, *18*).

To investigate this, we estimated RSN activity by measuring the proportion of variance explained by RSNs in the blood-oxygen-level-dependent (BOLD) signal from resting-state fMRI data. We analyzed four samples: 1) a large sample of young adults from the general population (*n* = 892), 2) a clinical sample with subacute back pain (SABP; *n* = 76, pain for 7-12 weeks), 3) a clinical sample with chronic back pain (CBP; *n* = 30, pain for more than 14 weeks), and 4) a clinical sample with treatment-resistant depression (*n* = 24), included to assess the specificity of our findings to pain. Both SABP and CBP groups were included due to known qualitative differences between these two phases of pain (*11*). Then, we applied Pareto optimality theory, which has been used to explain how evolutionary pressures and environmental demands shape populations through trade-offs across multiple tasks (*19–21*). Pareto optimality theory was applied using the Pareto task inference (ParTI) method (*22*); further details on both Pareto optimality theory and ParTI are provided in the Materials and Methods section.

The sample from the general population was used to define the neural archetypes, which were characterized using cognitive variables and enrichment analysis. The aim of the enrichment analysis was to infer the cognitive strategies optimized by each archetype. To explore the neuromodulatory systems involved, we compared the brain maps of these archetypes with the distribution of receptors and transporters for various neuromodulators, given their role in shaping both RSNs and behavioral traits (*23*). Then, we investigated the relationship between these archetypes and the clinical samples. By repeating the analyses *de novo* in the clinical pain samples, we replicated the archetypes, revealing connections between these archetypes and critical aspects of the pain experience.

## Results

### Study characteristics

To identify shared components across all participants, 19 RSNs were extracted using spatial independent component analysis (ICA) applied to the entire dataset, including all four samples (Fig. 1). For each participant, the proportion of variance explained by each RSN was expressed as a proportion of the total variance explained by all the 19 RSNs, which were supposed to capture most of the resting-state brain activity. To account for potential confounding factors, the effects of age, gender, and in-scanner movement (measured by three metrics, see Materials and Methods) were regressed out, resulting in an adjusted proportion of variance for each RSN. This adjusted metric was used in all subsequent analyses.

**Fig. 1.**
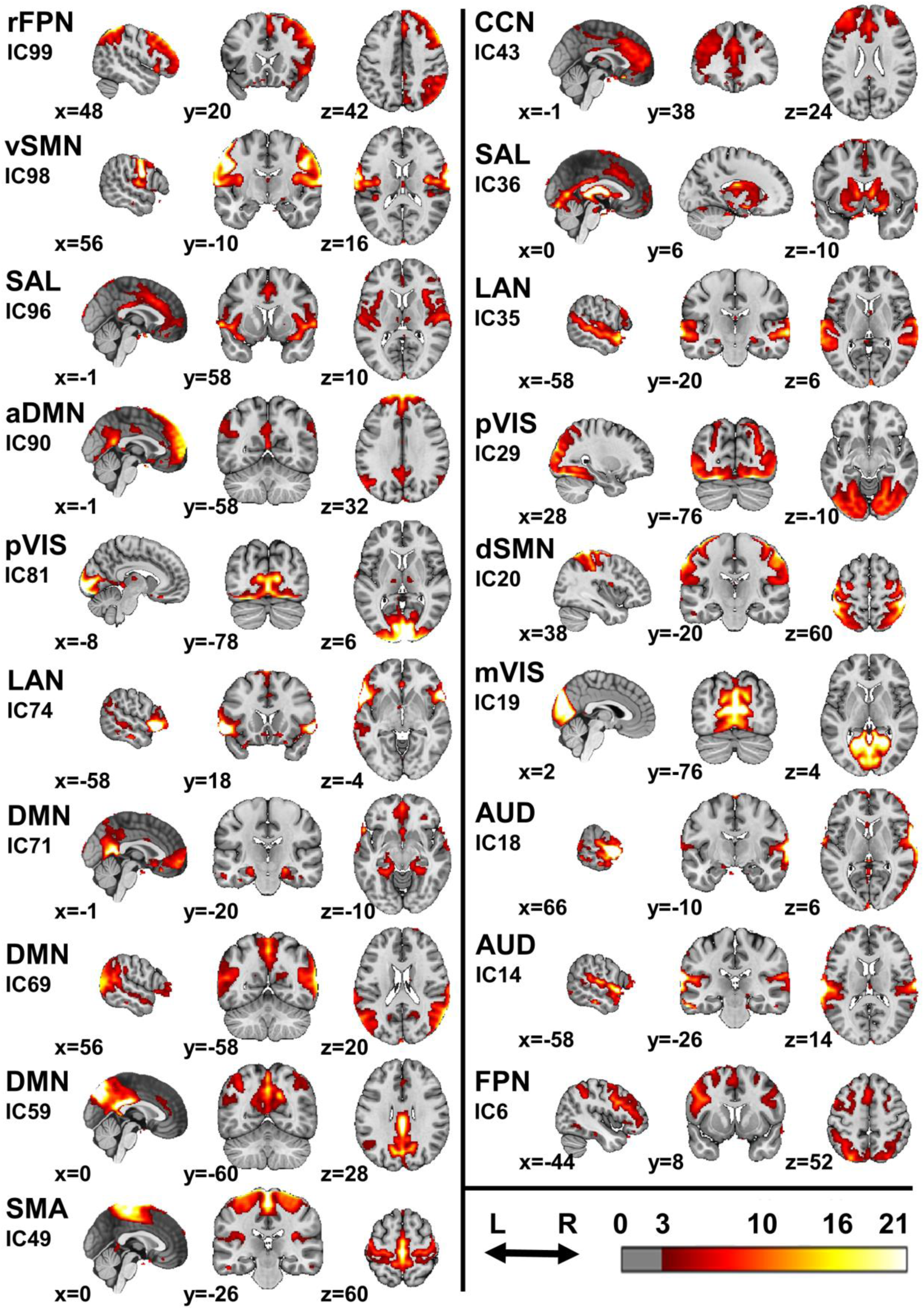
Resting-state networks. Aggregated images of the 19 RSNs derived from spatially independent component analysis. The color scale is expressed in arbitrary units, with a threshold of 3 applied alongside a cluster threshold of 10 voxels. RSNs are associated with their independent component numbers, which can be used to interpret the Supplementary Data. FPN = frontoparietal network; SMN = somatomotor network; SAL = salience network; DMN = default mode network; VIS = visual network; LAN = language network; SMA = supplementary motor area network; CCN = cognitive control network; r = right; d = dorsal; v = ventral; a = anterior; m = medial; p = peripheral; IC = independent component.

### The neural archetypes of the resting state optimize different cognitive strategies

Our primary analyses focused on the sample from the general population, as Pareto optimality theory predicts that only an evolutionarily adapted population should be confined within a polygon or polyhedron. The neural archetypes were defined using the adjusted proportion of variance of the 19 RSNs for each individual. While the neural archetypes were defined using only the RSNs, the enrichment analysis included also cognitive and behavioral features. In addition to the 19 RSNs, the enrichment analysis included questionnaires and behavioral tests assessing instrumental learning, decision-making abilities, temporal preferences for rewards, personality traits (impulsivity, sensation seeking, neuroticism, extraversion, openness, agreeableness, and conscientiousness), painful and non-painful somatic symptoms, depressive and anxiety symptoms, experiences of trauma, and average school grades (see Table S1 for a complete list). Confounders (age, gender, and in-scanner movement) that were previously regressed out from the proportion of variance of the RSNs were also included as features.

Plotting the first two principal components (PCs) of the adjusted proportion of variance (Fig. 2A) revealed that the sample was confined within a triangular shape defined by three archetypes (*t*-ratio test: P = 0.030). Enrichment analysis identified the RSNs and behavioral features significantly associated with each neural archetype (see Data S1 for results on all features). To visualize these findings (Fig. 2A), brain maps of the archetypes were constructed by calculating the weighted averages of the spatial ICs of the RSNs, with weights determined by effect sizes (rank biserial correlations; Data S1). These archetypal brain maps were compared to 44 independent brain maps from the neuromaps toolbox (*24*), mainly derived from positron emission tomography (PET) studies on brain receptors and transporters (Fig. 2B and Data S2). The three neural archetypes were named based on their optimized cognitive strategy: Archetype P (from *Punishment*), Archetype I (from *Impulsivity*), and Archetype C (from *Conscientiousness*).

**Fig. 2.**
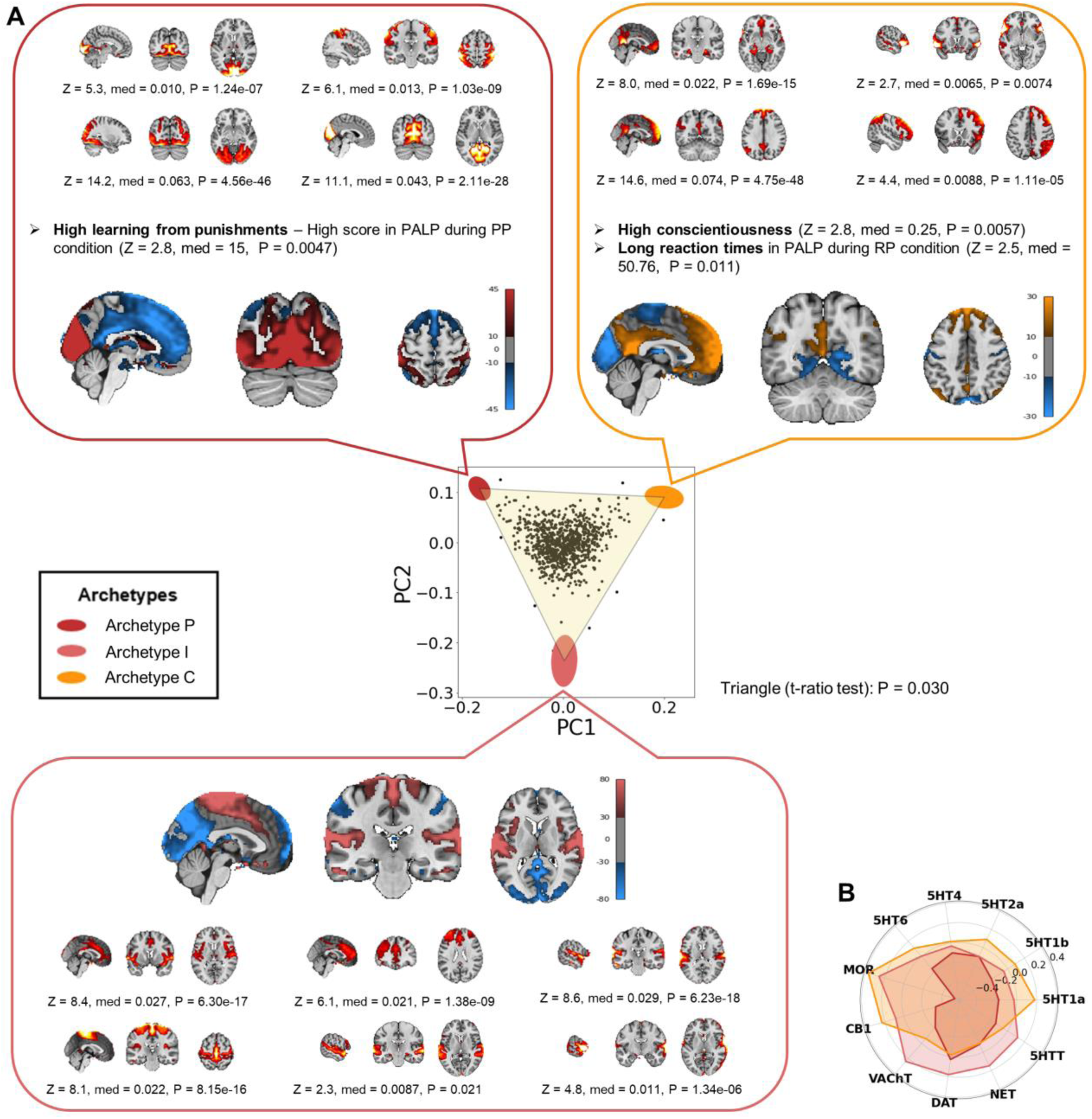
The neural archetypes of the resting state. **(A)** The first two principal components (PCs) of the adjusted proportion of variance explained by RSNs from the entire IMAGEN sample (*n* = 892) are shown. Colored ellipses represent the archetypes, with their centers marking the expected positions and their shapes indicating error margins, calculated via 10,000 bootstrap resampling. Brain maps of the neural archetypes are shown adjacent to each vertex. These maps were constructed by computing the weighted average of the spatial ICs of all the 19 RSNs, with weights corresponding to the effect sizes of each RSN. Enriched features are displayed alongside each archetype, including the RSNs that were maximally enriched, while minimally enriched RSNs are not individually shown (see Data S1 for all features). Color maps indicate the units of the brain maps for each archetype, as resulting from the weighted average of the spatial ICs. All tests of the enrichment analysis had 890 degrees of freedom. **(B)** Spider plot representing the similarity, expressed by Spearman’s correlations, between the brain maps of the archetypes and the distribution of several receptors and transporters (results for all the receptors and transporters are reported in Data S2). Z = z-score; med = median difference; P = P-value; PALP = Passive Avoidance Learning Paradigm; PP = punishment-punishment; RP = reward-punishment; MCQ = Monetary Choice Questionnaire; rho = Spearman’s rho; 5HTT = serotonin transporter; 5HT1a = serotonin receptor 1a; 5HT1b = serotonin receptor 1b; 5HT2a = serotonin receptor 2a; 5HT4 = serotonin receptor 4; 5HT6 = serotonin receptor 6; MOR = mu opioid receptor; CB1 = cannabinoid receptor 1; VAChT = vesicular acetylcholine transporter; DAT = dopamine transporter; NET = norepinephrine transporter.

**Archetype P** was optimized for punishment learning. It was maximally enriched in visual-somatosensory networks, including the medial and peripheral visual networks and a somatosensory network extending posteriorly into the dorsal attention network (Fig. 2A; e.g., peripheral visual network IC29, Z(890) = 14.2, P = 4.56 × 10^-46^). On the other hand, this archetype was minimally enriched in a group of RSNs including the anterior default mode network (DMN) and a salience network involving the anterior cingulate cortex (ACC) and insula (ACC-Insula salience network). It was negatively correlated with the brain maps of most neurotransmitter receptors and transporters, particularly the mu opioid receptor (MOR), the cannabinoid receptor 1 (CB1), and several serotonin receptors (Fig. 2B and Data S2). Behaviorally, this archetype was characterized by high learning from punishments, represented by higher scores in an instrumental learning task involving punishments (Fig. 2A; Z(890) = 2.8; P = 0.0047). The higher scores were due to fewer commission rather than omission errors (Data S1).

**Archetype I** may be optimized for specific aspects of impulsivity, including preference for immediate rewards and fast responses. It was enriched in several networks, including the ACC-Insula salience network; a cognitive control network involving the ACC and the dorsolateral prefrontal cortex (dlPFC); a somatomotor network involving somatomotor areas and the dorsal posterior insula; and auditory and language networks (Fig. 2A). Conversely, this archetype was minimally enriched in visual, somatosensory, and DMNs. Its brain pattern positively correlated with the distributions of acetylcholine, dopamine, and norepinephrine transporters, as well as MOR (Fig. 2B and Data S2). No behavioral feature was significant in the enrichment analysis of Archetype I, making it difficult to confidently infer its optimized strategy. We noticed that certain features associated with impulsivity approached statistical significance (Data S1). For instance, preference for immediate rewards, measured by the discounting rate of monetary rewards, was highest near Archetype I, while deliberation time in the Cambridge Gambling Task was lowest. These findings align with characteristics of impulsive behavior. However, contrary to expectations, both risk-taking and the overall proportion bet in the same task were also minimal near Archetype I. To further explore the cognitive strategies potentially optimized by Archetype I, we correlated Euclidean distances from this archetype with all the cognitive and behavioral measures (Data S1), and the two most correlated features were preference for immediate rewards (Spearman’s ρ(890) = −0.088, P = 0.0098) and motor impulsivity (Spearman’s ρ(890) = −0.081, P = 0.023). This pattern of RSNs and cognitive-behavioral features suggests that Archetype I may be optimized for specific aspects of impulsivity, including preference for immediate rewards and fast responses.

**Archetype C** was optimized for conscientiousness and self-control. It was enriched in the main and anterior DMNs, the right frontoparietal network (FPN), and the language network (Fig. 2A; e.g., anterior DMN IC90, Z(890) = 14.6, P = 4.75 × 10^-48^). At the same time, this archetype was minimally enriched in sensory networks (visual, somatosensory, and auditory). Its brain pattern was positively correlated with the brain maps of several receptors, such as MOR and CB1 (Fig. 2B and Data S2). Archetype C was behaviorally characterized by high conscientiousness (Z(890) = 2.8, P = 0.0057), a personality trait involving self-control, orderliness, and industriousness. Archetype C was also associated with long reaction times during an instrumental learning task involving both rewards and punishments (Fig. 2A; Z(890) = 2.5, P = 0.011).

### Chronic pain is linked to Archetype P

After defining the archetypes for an adapted sample, we examined their relationships with three clinical samples (SABP, CBP, and Depression). The clinical samples were projected onto the 2D space where the three archetypes were defined (Fig. 3A). Euclidean distances from the archetypes in the original 19D space significantly differed between the different samples (Kruskal-Wallis tests: Archetype P: χ^2^(3) = 76.0, P = 2.23 × 10^-16^; Archetype I: χ^2^(3) = 34.6, P = 1.49 × 10^-7^; Archetype C: χ^2^(3) = 36.9, P = 4.88 × 10^-8^). Both the SABP and CBP samples were significantly closer to Archetype P and significantly further from Archetypes I and C compared to the sample from the general population (Fig. 3B). Additionally, the CBP sample was significantly further from Archetype I than the SABP sample. Distances from the archetypes did not differ significantly based on medication (Table S2). Conversely, the Depression sample was not statistically different from the sample from the general population (Fig. 3B). To rule out potential site effects, we extracted the same ICs from a sample of healthy controls (*n* = 39) acquired alongside the clinical pain samples. These data were processed identically to the other datasets. The healthy controls were comparable to the general population. Relative to the healthy controls, the clinical pain samples were significantly closer to Archetype P and significantly further from Archetypes I and C (Fig. S1). Finally, while our primary focus was on the three main archetypes, the analyses were repeated with four archetypes, revealing a similar pattern of results (Figs. S2-S3 and Data S1).

**Fig. 3.**
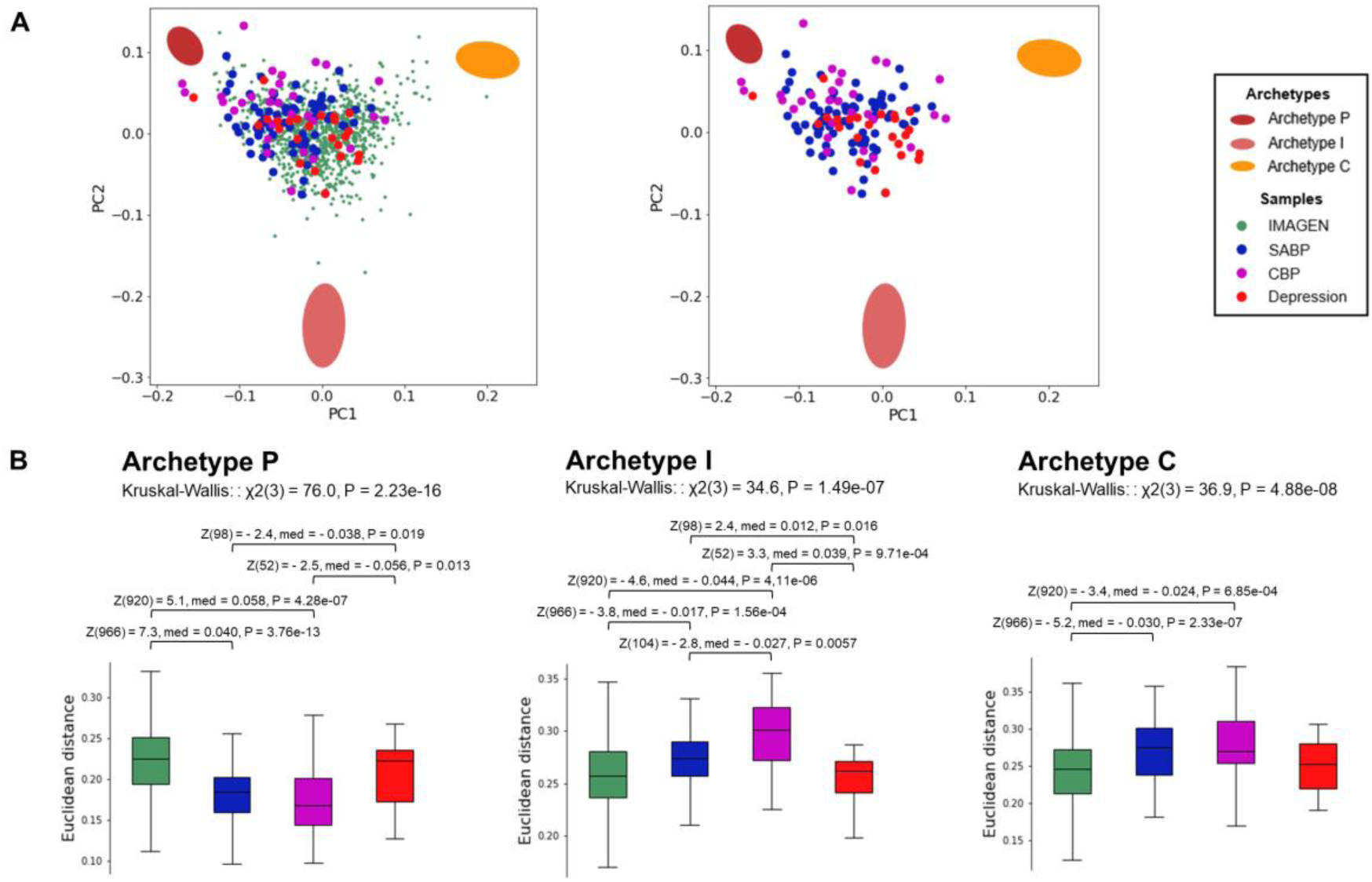
Clinical pain samples cluster near Archetype P. **(A)** The three clinical samples are projected onto the space defined by the first two principal components of the sample from the general population (IMAGEN), where the three archetypes were identified. The left plot includes both the IMAGEN and clinical samples, while the right plot displays only the clinical samples. **(B)** Euclidean distances of the clinical samples from the archetypes were analyzed using Kruskal-Wallis tests, followed by post-hoc Mann-Whitney U-tests (false discovery rate (FDR) < 0.05). SABP = subacute back pain; CBP = chronic back pain; χ2 = chi-squared; Z = z-score; med = median difference; P = p-value.

### Replication of neural archetypes and their relevance to the pain experience

As previously described, the neural archetypes presented above were computed using the sample from the general population. To replicate them in an independent sample, we recomputed the archetypes by applying the ParTI method *de novo* to the clinical pain sample, combining the SABP and CBP groups. Additionally, this approach ensured that the presence of chronic pain did not alter the fundamental archetype structure. While the neural archetypes were still defined using the 19 RSNs, the enrichment analysis included different cognitive and behavioral features than those used in the previous analysis based on the sample from the general population. Many of these features were related to key aspects of the pain experience, such as pain severity, interference, affective distress, support, life control, reactions from spouses or significant others, everyday activities, active coping, and catastrophizing. Additional features of the enrichment analysis included the adjusted proportion of variance of the 19 RSNs, education, and anxiety and depressive symptoms (see Table S3 for a complete list). Confounders such as age, gender, and in-scanner movement were also included as controls.

The pain samples significantly fell into a triangle (Fig. 4A; *t*-ratio test: P = 0.0028). The neural archetypes identified in this analysis closely mirrored those found in the sample from the general population, as evidenced by the similar brain maps of the archetypes (Fig. 4A) and the RSNs significantly enriched at each archetype (see Data S3 for all the features). This indicated that the fundamental archetype structure was not changed by chronic pain. Behaviorally, Archetype I was associated with high levels of catastrophizing (Z(104) = 2.6, P = 0.0093).

**Fig. 4.**
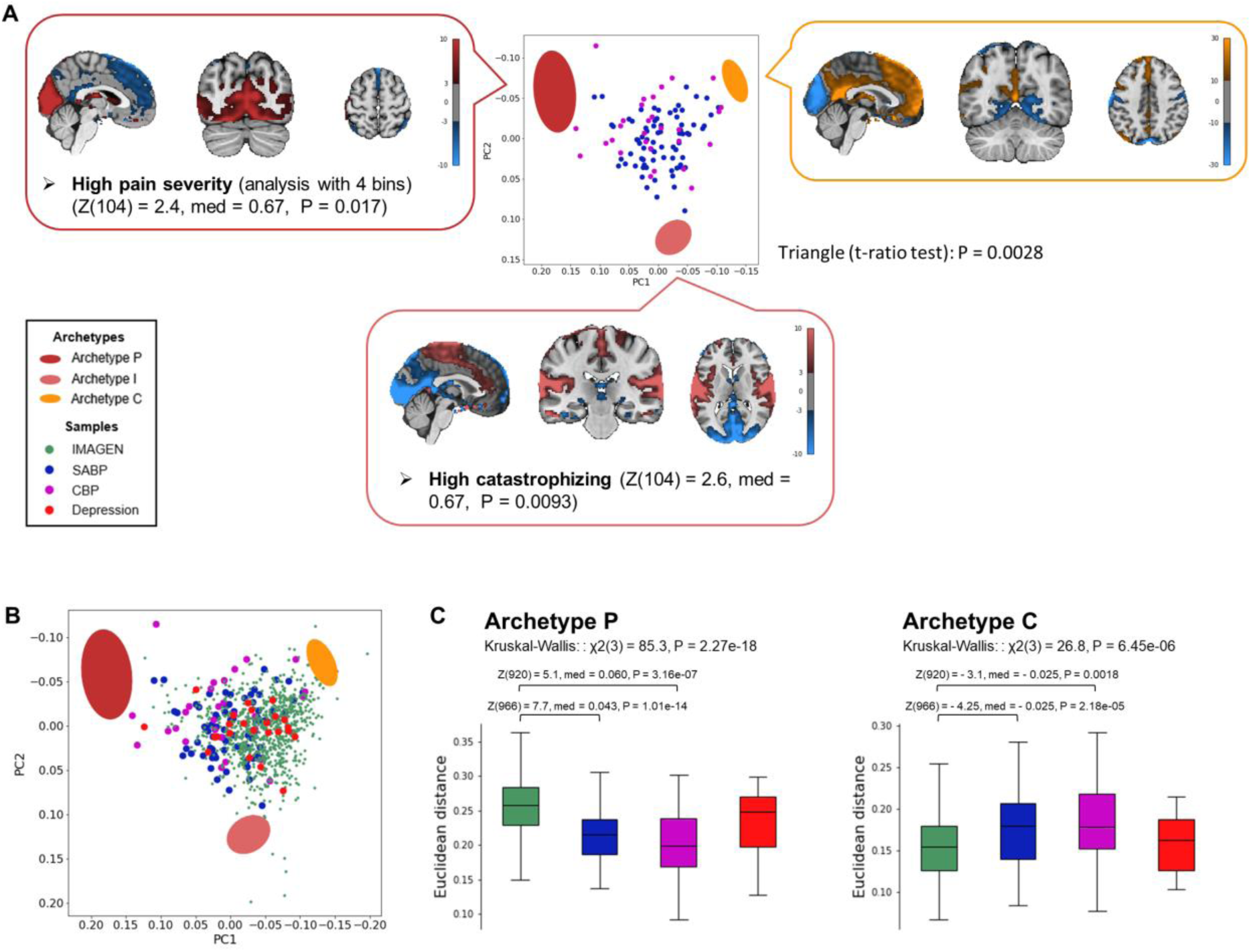
The three archetypes are independent of the sample used to define them. **(A)** The first two principal components (PCs) of the adjusted proportion of variance explained by RSNs from the pain sample combining individuals with SABP and CBP (*n* = 106) are shown. Colored ellipses represent the archetypes, with their centers corresponding to the expected positions and their shapes indicating error margins, calculated via 10,000 bootstrap resampling. The brain maps of the neural archetypes were constructed by computing the weighted average of the spatial ICs, with weights corresponding to the effect sizes of each RSN. Enriched features are displayed alongside each archetype. Color maps indicate the units of the brain maps for each archetype, based on the weighted average of the spatial ICs. **(B)** The sample from the general population (IMAGEN) and the Depression sample are projected onto the space defined by the first two principal components of the pain sample, where the three archetypes were replicated. **(C)** Euclidean distances from the archetypes of the different samples were analyzed using Kruskal-Wallis tests, followed by post-hoc Mann-Whitney U-tests (FDR < 0.05). Archetype I is not displayed, as it was not significant in the Kruskal-Wallis test. Z = z-score; med = median difference; P = P-value; aMCC = anterior midcingulate cortex.

Features that decay more slowly with the distance from the archetypes can be captured by dividing the sample into fewer bins during the enrichment analysis (*22*) (see Materials and Methods). When using four bins in the enrichment analysis, Archetype P was associated with high pain severity (Fig. 4A and Data S3 for all features; Z(104) = 2.4, P = 0.017). Then, we projected the sample from the general population and the Depression sample onto the 2D space where the three archetypes based on the pain samples were defined (Fig. 4B). Euclidean distances from the archetypes in the original 19D space significantly differed between the different samples for Archetypes P and C, but not I (Kruskal-Wallis tests: Archetype P: χ^2^(3) = 85.3, P = 2.27 × 10^-18^; Archetype I: χ^2^(3) = 6.7, P = 0.08; Archetype C: χ^2^(3) = 26.8, P = 6.45 × 10^-6^). Again, both the SABP and CBP samples were significantly closer to Archetype P and significantly further from Archetype C compared to the sample from the general population (Fig. 4C). This findings indicated that the archetypes are independent from the sample used to define them, and that the distance from the archetypes is a neural-cognitive trait relevant for chronic pain.

## Discussion

In this study, we showed that individuals’ RSN patterns result from trade-offs between extreme neural types, called archetypes, optimized for specific cognitive strategies. These RSN patterns were quantified by the proportion of variance of the BOLD signal explained by the RSNs. To further characterize the neural archetypes, we compared their brain maps with the distribution of receptors and transporters for various neuromodulators. Additionally, we investigated the relationship of these archetypes with three clinical samples, showing that people with SABP and CBP, compared to individuals from the general population, are significantly closer to Archetype P, characterized by high learning from punishments. Finally, we replicated the neural archetypes in the pain sample, finding connections between the archetypes and critical aspects of the pain experience.

### The neural archetypes of the resting state optimize different cognitive strategies

First, we aim to interpret our findings in relation to the cognitive strategies that are prioritized by individuals closer to each archetype. We will then use this interpretation to explore how these strategies may contribute to the development of chronic pain. We attempted to infer the optimized strategies by finding the RSNs and cognitive-behavioral features enriched at each archetype. Archetype P was maximally enriched in visual-somatosensory networks and minimally enriched in DMNs and the ACC-Insula salience network. Behaviorally, Archetype P was characterized by high learning from punishments, represented by higher scores in an instrumental learning task involving punishments, which were driven by fewer commission errors. In other words, when individuals close to Archetype P made a mistake and were punished, they quickly adjusted their behavior to avoid repeating that mistake. They made fewer impulsive errors, which led to higher overall scores. The brain map of Archetype P was negatively correlated with brain maps of the mu opioid receptor, the cannabinoid receptor 1, the serotonin transporter and all the serotonin receptors we tested. Serotonin plays an important role in behavioral inhibition, particularly in response to aversive stimuli and contexts, and modulates large-scale functional connectivity changes (*23*, *25–29*). For instance, the serotoninergic system modulates the DMN in complex ways, as shown by studies that pharmacologically targeted serotonin receptors and transporters (*30–32*). The cannabinoid receptor 1 is critically involved in the extinction of aversive memories, and its deficiency promotes passive coping strategies in threatening situations (*33*). Mu opioid receptor agonists reduce the sensitivity to aversive stimuli and inhibit fear conditioning (*34*). Overall, Archetype P may optimize learning from punishments through increased sensitivity to aversive stimuli and enhanced self-control in aversive situations.

Archetype I was associated with several RSNs, including the ACC-Insula salience network and a somatomotor network. We could not confidently infer any cognitive strategy associated with this archetype, as no cognitive feature was significant in the enrichment analysis. However, additional analyses suggested a potential link between Archetype I and specific aspects of impulsivity, including preference for immediate rewards and fast reponses. Supporting this idea, the salience network plays a key role in processing relevant stimuli, including rewards (*35*), and the functional connectivity between the right anterior insula and the ACC has been negatively correlated with monetary discounting (*36*). Interestingly, prioritizing immediate rewards may represent the most adaptive strategy under stressful conditions, where the future is uncertain and cannot be relied upon (*37*, *38*). This possible association between Archetype I and impulsivity is particularly intriguing given a previous study (*n* = 1206) which identified three distinct cognitive archetypes based on time preferences for rewards: one favoring smaller immediate rewards, another preferring larger delayed rewards, and a third adopting a more flexible strategy (*39*). These cognitive archetypes were associated with differences in brain structure and functional brain connectivity. However, we argue that archetypes should be defined based on functional brain activity rather than cognitive traits, as neural circuits are optimized for cognitive functions, not the reverse.

Archetype C was characterized by the DMN, the right FPN, and the language network, while being minimally enriched in sensory networks. Behaviorally, Archetype C was associated with high conscientiousness and long reaction times in an instrumental learning task involving both rewards and punishments, suggesting a more careful decision-making. Conscientiousness is a personality trait encompassing self-control, industriousness, orderliness, responsibility, and adherence to rules (*40*). The neural and behavioral features of Archetype C form a cohesive pattern. The DMN is associated with self-referential thinking, mind-wandering, episodic memory, semantic memory, and social cognition, and it is typically suppressed during externally driven tasks (*16*). The FPN is associated with executive control, managing top-down, goal-directed processes while flexibly modulating other brain networks (*41*). Like the other archetypes, Archetype C likely involves individual variations in both the dopaminergic and serotonergic systems. DA is integral to exertion of effort, sustained task engagement, goal-directed behavior, and instrumental learning from rewards and punishments (*42–44*). Serotonin contributes to behavioral inhibition and the processing of verbal memory (*25*). In summary, the neural and cognitive characteristics of Archetype C suggest it embodies a future-oriented, goal-directed cognitive strategy optimized for self-control and conscientious decision-making.

The contrast between Archetype I and Archetype C – one potentially optimized for fast responses, the other for self-control – echoes the fast-and-slow thinking and dual process models that have shaped psychology, economics, and related disciplines (*45–47*). These models distinguish between two ways of thinking, commonly referred to as System 1 and System 2: the former is fast, intuitive, and effortless, while the latter is slow, deliberate, and effortful. Archetypes I and C may reflect these two cognitive systems, while Archetype P may represent a distinct strategy in which self-control and slow thinking are preferentially adopted in aversive situations.

In summary, the three archetypes point to the optimization of cognitive and behavioral strategies adapted to different environmental demands. These strategies are mainly based on differences in instrumental learning, impulsivity/behavioral inhibition, and decision-making. Neuromodulatory systems are involved in all these behaviors and can influence RSNs, suggesting that they may explain the connection between the neural and cognitive features of the archetypes, although limitations of our analysis, gaps in the literature and methodological differences across studies make these connections difficult to discern. Future research should investigate these archetypes in a longitudinal way, including genetic data to disentangle the relationship between genes, RSNs, and cognition.

### Chronic pain is linked to Archetype P

We hypothesized that mental disorders may be linked to extreme neural types that prioritize specific cognitive strategies. Our results showed that clinical pain samples mapped closer to Archetype P, and further from Archetypes I and C, than a sample from the general population. This pattern was not observed in a clinical sample with treatment-resistant depression, suggesting specificity to pain-related conditions. This indicates that chronic pain is associated with a cognitive strategy characterized by increased learning from punishments and enhanced behavioral inhibition in response to aversive stimuli, possibly mediated by the serotonergic system. Pain is an aversive stimulus and drives learning processes (*2–4*). Instrumental learning plays an important role in the development of chronic pain, as individuals often develop avoidance or escape behaviors (e.g., protective postures, limping) which can increase pain over time through reinforcement learning mechanisms (*2–7*). According to the motivation-decision model, pain is not a passive experience but rather the result of conflicting motivational systems (e.g., avoiding harm, pursuing rewards) constantly engaging in neural decision processes that either prevent or facilitate pain perception by controlling the descending modulatory pathway (*48*). Notably, this pathway inhibits pain by activating mu opioid receptors (*48*) and Archetype P was strongly negatively correlated with the brain-wide distribution of mu opioid receptors. Then, individuals close to Archetype P may prioritize harm avoidance over reward seeking, thereby facilitating behaviors that favor avoidance over engagement. This dynamic can sustain the chronic pain state, establish pain-enhancing avoidance behaviors, and reduce exposure to rewarding activities that might otherwise alleviate pain. Conversely, Archetypes I and C may protect against chronic pain by prioritizing immediate and future rewards, respectively.

The SABP sample was closer to Archetype I than the CBP sample, suggesting that chronic pain is associated with a reduced preference for immediate rewards. This aligns with research indicating that the subacute and chronic stages of pain involve distinct alterations in both the neural and cognitive aspects of reward processing (*9*, *11*, *49*). Specifically, people with SABP who later transition to CBP are more responsive to reward prediction errors (*11*), while CBP is associated with reduced reward responsiveness (*9*, *50*), consistent with the greater distance from Archetype I.

Two possibilities arise regarding the development of chronic pain: (1) individuals closer to Archetype P may be more predisposed to developing chronic pain (trait), or (2) they shift closer to Archetype P as the condition progresses (state). More broadly, this raises the open question of whether individuals can move within the triangle over time and what the timescale of such changes might be. We would expect the position of an individual within the triangle to be relatively stable, reflecting the distinctive strategies they employ. Furthermore, age was controlled for in our analysis, and no significant associations with any archetype were observed. However, addressing these questions would require a longitudinal study.

### Replication of neural archetypes and their relevance to the pain experience

The three neural archetypes found in the sample from the general population were replicated in an independent sample of individuals with SABP and CBP, showing their relevance to the pain experience. Pain severity was higher near Archetype P, aligning with the motivation-decision model of pain described above, the evidence that aversive pain memories can amplify pain perception, and the effects of short-term avoidance of pain, which establishes pain behaviors such as protective postures that increase pain over time (*2–4*, *7*, *48*, *51*). Supporting the hypothesis that individuals closer to Archetype P may be more sensitive to pain, a study showed that lowered resting-state fMRI signal variability in the DMN, somatosensory cortex, and salience network was linked to greater pain sensitivity, as measured by temporal summation of pain (*52*). Additionally, pain catastrophizing was more prominent near Archetype I, consistent with studies linking a preference for immediate rewards to anxiety (*39*, *53*). In fact, pain catastrophizing involves excessive worry about pain and positively correlates with anxiety (*54*).

Further replicating the previous results, we showed that clinical pain samples mapped closer to Archetype P and further from Archetype C than the sample from the general population. Importantly, these findings revealed that independently of the sample used to define the archetypes, the distance of an individual from the archetypes is a neural-cognitive trait relevant for chronic pain.

### Limitations

This study has several limitations. First, we had a limited set of cognitive and behavioral features, and other constructs may be critical to better understand the tasks optimized by the archetypes. Second, the age differences among our samples may have influenced the results, as research has shown variations in associative learning and impulsivity between younger and older individuals (*55–59*). However, age was regressed out in our analyses and found not to be significantly associated with any archetype. Additionally, we found similar neural archetypes independent of the age distribution of the sample used to define them. Third, missing values in questionnaires and behavioral tests may have affected the results. In particular, catastrophizing had a high percentage of missing values (25%), while all the other features significantly associated with the archetypes had a low percentage of missing values (< 5%).

## Conclusion

In this study, we showed that individual differences in RSN patterns correspond to distinct cognitive strategies, consistent with an evolutionary perspective in which individuals navigate trade-offs between optimizing multiple cognitive functions. These strategies were particularly relevant to chronic pain, as individuals with chronic pain tended to be close to the archetype optimized for punishment learning, while being distant from those associated with self-control or impulsivity.

More broadly, we introduced a novel framework for studying clinical populations by contextualizing them in relation to the general population. This framework conceptualizes neural-cognitive traits as distances from extreme forms, called archetypes, optimized for different functions. Future research should investigate how individuals change over time in relation to the archetypes, the role of genetic influences, and the connections between these archetypes and other disorders. The findings presented here provide insights into both clinical disorders and human behavior, paving the way for new questions and directions in research.

## Materials and Methods

### Datasets

#### IMAGEN

IMAGEN is a longitudinal dataset (https://imagen-project.org/) collected in eight centers in Europe (London, Nottingham, Dublin, Paris, Berlin, Dresden, Hamburg, Mannheim). Written informed consent was obtained from all participants by the IMAGEN consortium, and the study received approval from the following ethics committees: the institutional ethics committee of King’s College London (PNM/10/11-126), University of Nottingham (D/11/2007), Trinity College Dublin (SPREC092007-01), Technische Universitat Dresden (EK 235092007), Commissariat a l’Energie Atomique et aux Energies Alternatives, INSERM (2007-A00778-45), University Medical Center at the University of Hamburg (M-191/07) and at the medical ethics committee of the University of Heidelberg (2007-024N-MA) in Germany. The study was conducted in accordance with the Declaration of Helsinki. To facilitate comparisons with the other datasets, we used resting-state fMRI data from the second follow-up phase, when participants were already adult. The initial sample size included 946 individuals. Participants with low-quality data were excluded based on both visual inspection after preprocessing and an assessment of the proportion of variance explained by noise ICs. Additionally, participants with more than 50% volume outliers were excluded, leaving a final sample of 892 participants (mean age: 19.0 ± 0.7 years, 55% female). Demographic data for the participants of all centers are provided in Table S4. Probabilities for diagnoses of mental disorders are reported in Table S5. Additional details about the IMAGEN dataset are available in its documentation (https://imagen-project.org/) and related publications (https://imagen-project.org/?page_id=471).

#### CBP-PREDICT

CBP-PREDICT is a longitudinal dataset that includes individuals with SABP and CBP. Data were collected at the Central Institute of Mental Health in Mannheim as part of the Heidelberg Pain Consortium (SFB1158, https://www.sfb1158.de/). All procedures received approval from the Ethics Committee of the Medical Faculty Mannheim of Heidelberg University and adhered to the principles of the Declaration of Helsinki. Using the same quality control criteria applied to the IMAGEN dataset, the final sample sizes were 76 for SABP (mean age: 35.5 ± 13.4 years, 71% female) and 30 for CBP (mean age: 41.4 ± 16.1 years, 53% female).

#### IAdapt

The IAdapt dataset contains a clinical sample of outpatients with treatment-resistant depression. It was collected as part of a single-blind, randomized, crossover, exploratory trial conducted at the Noninvasive Neuromodulation Centre (CEMNIS) in Strasbourg (NCT02863380) (*60*). The study aimed to compare different neuromodulation procedures and was conducted in accordance with the principles of the Declaration of Helsinki (World Medical Association, 2013), with approval from the local ethics committee (CPP Est IV). All patients provided informed consent prior to participation. Resting-state fMRI data were analyzed at baseline, before the start of neuromodulation therapy. Using the same quality control criteria applied to the IMAGEN dataset, all 24 participants had high-quality data and were included in the analyses (mean age: 54.6 ± 13.3 years, 71 % female).

### Functional magnetic resonance imaging: data acquisition

#### IMAGEN

Scanning parameters were standardized across the eight centers participating in IMAGEN, despite using different 3 Tesla scanners (Siemens, Munich, Germany; Philips, Best, The Netherlands; GE Healthcare, Chicago, USA). The resting-state fMRI protocol included a repetition time (TR) of 2.2 seconds, an echo time (TE) of 30 milliseconds, and a resolution of 64 × 64 pixels. A gradient-echo echo-planar imaging (EPI) T2*-weighted sequence was employed to acquire 40 slices in a sequential descending order, with a slice thickness of 2.4 mm and a 1 mm gap, using an acceleration factor of 2. Detailed imaging protocols from all centers are available on https://github.com/imagen2/imagen_mri. Scanning parameters for all the centers are reported in Table S6 for functional data and Table S7 for anatomical data.

#### CBP-PREDICT

Data were acquired using a 3 Tesla Tim TRIO whole-body scanner (Siemens Healthineers, Erlangen, Germany). The protocol used a T2*-weighted gradient-echo EPI sequence with GRAPPA technique (TR = 2.1 s, TE = 23 ms, matrix size = 96 × 96, acceleration factor = 2, field of view (FoV) = 220 x 220 mm^2^, flip angle (α)= 90°). Thirty-six contiguous axial slices were acquired, each 3 mm thick, with an in-plane voxel size of 2.3 × 2.3 mm and no inter-slice gap. The structural MRI protocol used a T1-weighted magnetization prepared rapid gradient echo (MPRAGE) sequence with 192 sagittal slices and voxel dimensions of 1 x 1 x 1 mm^3^ (TR = 2.3 s, TE = 2.98 ms, matrix size = 240 × 256 pixels, FoV = 240 × 256 mm², flip angle (α)= 9°).

#### IAdapt

Imaging was conducted on a 3 Tesla MAGNETOM Verio scanner with syngo MR B17 software (Siemens Healthineers, Erlangen, Germany). The protocol employed a T2*-weighted gradient-echo EPI sequence with a multiband acceleration factor of 8. Forty contiguous axial slices were acquired with voxel dimensions of 3 × 3 × 3 mm (TR = 407 ms, TE = 36.6 ms, matrix size = 64 × 64 pixels, FoV = 192 × 192 mm², flip angle (α)= 40°). The structural MRI protocol used a T1-weighted MPRAGE sequence with 224 sagittal slices and voxel dimensions of 0.7 x 0.7 x 0.7 mm^3^ (TR = 2.4 s, TE = 2.41 ms, matrix size = 320 × 320 pixels, FoV = 224 × 224 mm², flip angle (α)= 8°).

### Functional magnetic resonance imaging: data processing

The same preprocessing was applied to all datasets using fMRIPrep 21.0.1 (*61*), an automatic preprocessing pipeline based on Nipype 1.6.1, with two exceptions. First, B0 distortion correction was performed only for the IMAGEN dataset, and only when a corresponding field map could be successfully applied. This was possible for 632 out of 892 participants (71%). To ensure that this was not influencing the results, we replicated most findings using only the participants without a field map (Fig. S4 and Data S4). Notably, this time also the sample with treatment-resistant depression was significantly closer to Archetype P than the sample from the general population. Second, slice-time correction was not required for the IAdapt dataset due to its short repetition time (TR = 407 ms), which was negligible compared to the hemodynamic response. A detailed description of the preprocessing steps executed by fMRIPrep is provided in the Supplementary Methods. The output reports of fMRIPrep were visually inspected to exclude low-quality data.

Subsequently, data were denoised by adding eight physiological regressors to the matrix composed of both the noise and signal components calculated by an ICA-based strategy for automatic removal of motion artifacts (ICA-AROMA), a step of the fMRIPrep pipeline designed to efficiently remove artifacts from the signal. The eight physiological regressors included the mean signals from white matter and cerebrospinal fluid, along with their first derivatives, squares, and squared derivatives. Denoising was performed using the *fsl_regfilt* function in FSL 6.0.

To harmonize data across all three datasets, the scanning duration for each participant was aligned to the shortest duration observed across the datasets (6 minutes), resulting in varying numbers of volumes per participant. This adjustment was intended to capture psychological dynamics over comparable timeframes and address biases related to increasing sleepiness during resting-state scans, as well as a recently discovered non-neuronal physiological effect involving time-dependent inflation of brain-wide functional connectivity (*62*).

The data were spatially smoothed using a 4 mm full-width half-maximum Gaussian kernel using the *fslmaths* function in FSL 6.0. Outlier volumes flagged by fMRIPrep were replaced with random values drawn from a Gaussian distribution matching the mean and standard deviation of the original time course, calculated after excluding the outlier volumes. Volumes were classified as outliers based on thresholds of mean framewise displacement (mFD > 0.5 mm) and mean standardized spatial root mean square after temporal differencing (mDVARS > 1.5), as detailed in Supplementary Methods.

### Functional magnetic resonance imaging: group ICA

To obtain the same RSNs across participants and samples, we applied group spatial ICA to the combined dataset using the group ICA of fMRI toolbox (GIFT) 4.0.5.0 (*63*) in MATLAB R2024a. The Infomax algorithm was used to estimate 100 ICs (*64*). The default settings in GIFT were adjusted as follows: 1) A brain mask covering the minimal common brain area across all samples was applied (see Data availability), 2) Intensity normalization was enabled, 3) Data dimensionality was reduced in a single step using group-level principal component analysis with unstacked multi-power iteration (*65*). This approach was chosen as it was computationally feasible given the size of the combined dataset. All other parameters were left at their default settings, and no additional scaling was applied to the resulting ICs.

### Adjusted proportion of variance explained by each RSN

The aggregate images of the 100 ICs were visually inspected to identify and exclude noise-related components. This process resulted in the identification of 19 RSNs (Fig. 1). For each RSN, we calculated the proportion of explained variance by dividing its variance by the total variance summed across all 19 RSNs. This normalization ensured comparability across different samples by accounting for noise-related variability. To further refine the analysis, we adjusted the proportion of variance by regressing out the effects of age, gender, and in-scanner movement. In-scanner movement was quantified using three metrics: mFD, mDVARS, and the percentage of outlier volumes. These adjustments provided an estimate of the contributions of the networks to each participant’s resting state, controlling for confounding factors.

### Behavioral data

#### IMAGEN

Data were obtained from psychological questionnaires and behavioral tests from the IMAGEN dataset reported in Table S1. This table also details the specific constructs included in the analyses and the percentage of missing values, which were imputed using the median. These measures evaluated a broad range of constructs, including personality traits (neuroticism, extraversion, openness, conscientiousness, agreeableness), impulsivity (attention, motor, planning), temporal preferences for rewards, sensation seeking, instrumental learning (involving punishments and/or rewards), decision-making abilities, painful and non-painful somatic symptoms, depressive and anxiety symptoms, experiences of trauma, and average school grades.

#### CBP-PREDICT

Data were obtained from the psychological questionnaires from the CBP-PREDICT dataset reported in Table S3, which also shows the percentage of missing values, imputed using the median. These scales assessed the following constructs: pain severity, interference, affective distress, support, life control, responses by a significant other (punishing, solicitous, and distracting responses), activities away from home and social activities, household chores, outdoor work, helplessness, resourcefulness, active coping, catastrophizing, depressive symptoms, and anxiety symptoms.

### Pareto optimality theory

Pareto optimality theory provides a framework to understand multitask optimization problems. Systems that need to optimize multiple tasks simultaneously face inherent trade-offs when it is impossible to maximize performance in all tasks at once. The solution to these problems is not unique, but rather consists of the set of all optimal trade-offs. This concept has been applied to explain how evolutionary processes shape the traits observed in biological systems (*19–21*).

In this framework, the space of traits provides a geometrical representation of all possible configurations of traits. A specific trait may enhance performance in one task but reduce it in another. The net effect of a trait on overall fitness depends on the relative importance of each task in a given environment. The traits observed in an evolutionarily adapted population represent the set of optimal trade-offs. These traits are combinations, or weighted averages, of specialized forms called archetypes, each of which being the theoretical extreme phenotype that would be specialized in a single task. The performance in the optimized task decreases with the distance in trait space from the corresponding archetype.

When a population is evolutionarily adapted and reaches Pareto optimality, the traits of its individuals are confined within a geometric shape in trait space - a polygon or polyhedron whose vertices represent the archetypes. The dimensionality of this shape is determined by the number of optimized tasks: with two tasks, traits align along a line connecting two archetypes; with three, they form a triangular region; with four, they occupy a tetrahedron in the trait space. Observing traits constrained within such a structure suggests that they have been positively selected, while those falling outside have likely been selected against, indicating maladaptation. This geometrical representation provides a way to visualize how evolutionary pressures and environmental constraints shape populations through trade-offs across multiple tasks.

Pareto optimality theory has been applied to explain morphological traits in animals (*19*, *66*), the evolution of ammonite shells after mass extinctions (*67*), gene expression patterns in bacteria and human cells (*19*, *68*, *69*), the structure of polymorphisms (*20*), *Escherichia coli* proteome (*70*), animal behavior (*71*), and time preferences for reward in humans (*39*).

### Pareto task inference (ParTI) method

The ParTI method was implemented using the MATLAB software package provided by the Uri Alon Lab (https://www.weizmann.ac.il/mcb/alon/download/pareto-task-inference-parti-method) (*22*). All steps described in this and the next section (Enrichment analysis) were executed within this software package, with few modifications of the source code (see Code availability).

The ParTI analysis followed four key steps: (1) performing principal component analysis, (2) estimating the positions of the archetypes, (3) assessing the statistical significance of the best-fit polygon or polyhedron, and (4) conducting enrichment analysis to identify features associated with each archetype.

The adjusted proportion of variance of the 19 RSNs from all participants in the IMAGEN sample served as input for the ParTI analysis. First, a PCA was performed, centering the data to have a mean of zero without standardization. The data were then projected onto the first *n* – 1 PCs, where *n* represents the number of archetypes. The polygon or polyhedron defined by the archetypes was identified using the Simplex Identification via Split-and-Augmented Lagrangian (SISAL) algorithm (*72*). The uncertainty in archetype positions was quantified using a bootstrap procedure that resampled the data with replacement 10,000 times.

Statistical significance of the best-fit polygon/polyhedron was assessed using the t-ratio test, which compares the volume of the polygon/polyhedron to the volume of the convex hull of the data. This test was performed by generating 10,000 randomized datasets, calculating their t-ratios, and comparing these values to the original t-ratio to derive an empirical P-value.

### Enrichment analysis

Enrichment analysis is the last step of the ParTI method. In order to infer which tasks are optimized by the archetypes, the ParTI method determines which features are significantly enriched in the individuals closest to the archetypes. We modified the original code to also identify features that are minimally enriched (see Code availability).

The procedure begins by sorting features based on the Euclidean distance from the archetype of each individual and dividing the features into ten bins (default). For continuous features, the bin closest to the archetype is compared to the rest of the data using the Mann-Whitney U-test. Here, we implemented the calculation of the rank-biserial correlation coefficient derived from the U statistics (*73*), which was used to produce the brain maps of the archetypes as described in the Result section. The brain maps of the archetypes were compared to 44 molecular and functional brain maps from the neuromaps toolbox (*24*), using Spearman’s rho correlation coefficients to assess the similarity of their brain maps and testing this similarity through 10,000 autocorrelation-preserving null models. For discrete features, the data are first transformed into Boolean variables, after which the significance of the bin closest to the archetype is assessed using the hypergeometric test. For both feature types, false discovery rate (FDR) correction is applied with the default significance threshold of 0.1 (*22*).

To mitigate potential circularity arising from using RSNs both to define archetypes and in the enrichment analysis, a leave-one-out procedure was performed. In this process, one RSN is excluded in each iteration, the positions of the archetypes are recalculated, and the enrichment analysis is repeated. This ensures that the features identified as significantly enriched near the archetypes remain robust even when individual RSNs are omitted. Given that some RSNs belong to the same functional systems (e.g., DMN), we conducted the enrichment analysis in two additional ways to account for system-level redundancy. First, we used RSNs as individual features and removed entire RSN systems during the leave-one-out procedure (Data S5). Second, we averaged the RSNs within each system to create composite features and then removed entire RSN systems during the leave-one-out process (Data S6). These controls ensured that the enrichment analysis was not biased by overlapping or system-level contributions of RSNs.

## Author Contributions

Conceptualization: F.S., J.R.F., L.D.J., M.L., H.F.; Methodology: F.S., J.R.F., R.S., E.S., M.L., H.F.; Data processing and analyses: F.S.; Visualization and interpretation of results: F.S., L.D.J., J.R.F., M.L., H.F.; Supervision: J.R.F., H.F.; Writing – original draft: F.S.; Writing – review & editing: F.S., L.D.J., R.S., J.R.F., E.S., M.L., H.F.; Data contribution: IMAGEN; IMAGEN study design: T.B., A.L.W.B., R.B., S.D., H.G., P.G., A.G., A.H., J.L.M., M.L.P.M., E.A., D.P.O., L.P., M.N.S., S.H., N.H., N.V., H.W., R.W., G.S., F.N., H.F.

## IMAGEN Consortium

Tobias Banaschewski, Arun L. W. Bokde, Rüdiger Brühl, Sylvane Desrivières, Hugh Garavan, Penny Gowland, Antoine Grigis, Andreas Heinz, Jean-Luc Martinot, Marie-Laure Paillère Martinot, Eric Artiges, Dimitri Papadopoulos Orfanos, Luise Poustka, Michael N. Smolka, Sarah Hohmann, Nathalie Holz, Nilakshi Vaidya, Henrik Walter, Robert Whelan, Gunter Schumann, Frauke Nees & Herta Flor

## Data and materials availability

The datasets analyzed during the current study are available upon reasonable request from the corresponding author F. Scarlatti (CBP-PREDICT), authors L. Dormegny-Jeanjean and J. R. Foucher (IAdapt – as these data are considered sensitive, access can only be granted to named individuals and in compliance with ethical and legal procedures governing their reuse), or the IMAGEN consortium (IMAGEN dataset, see https://imagen-project.org/the-imagen-dataset/), respectively.

The full reproducible code can be found at the Open Science Framework (https://osf.io/kngfb/?view_only=0ef8e252435a42eebf60bfdf4b3c061b). Additionally, all data necessary to reproduce and build upon this work can be found at the same link, including the 100 ICs, the proportion of variance for each RSN for each participant, the positions of the archetypes, and the brain mask applied to all participants.

