## Supplementary Materials for "Chronic pain is linked to a resting-state neural archetype that optimizes learning from punishments"

**This PDF file includes:**

Figs. S1 to S4
Tables S1 to S7
Supplementary Methods
References

**Other Supplementary Materials for this manuscript include the following:**

Data S1 to S6

### Supplementary Figures

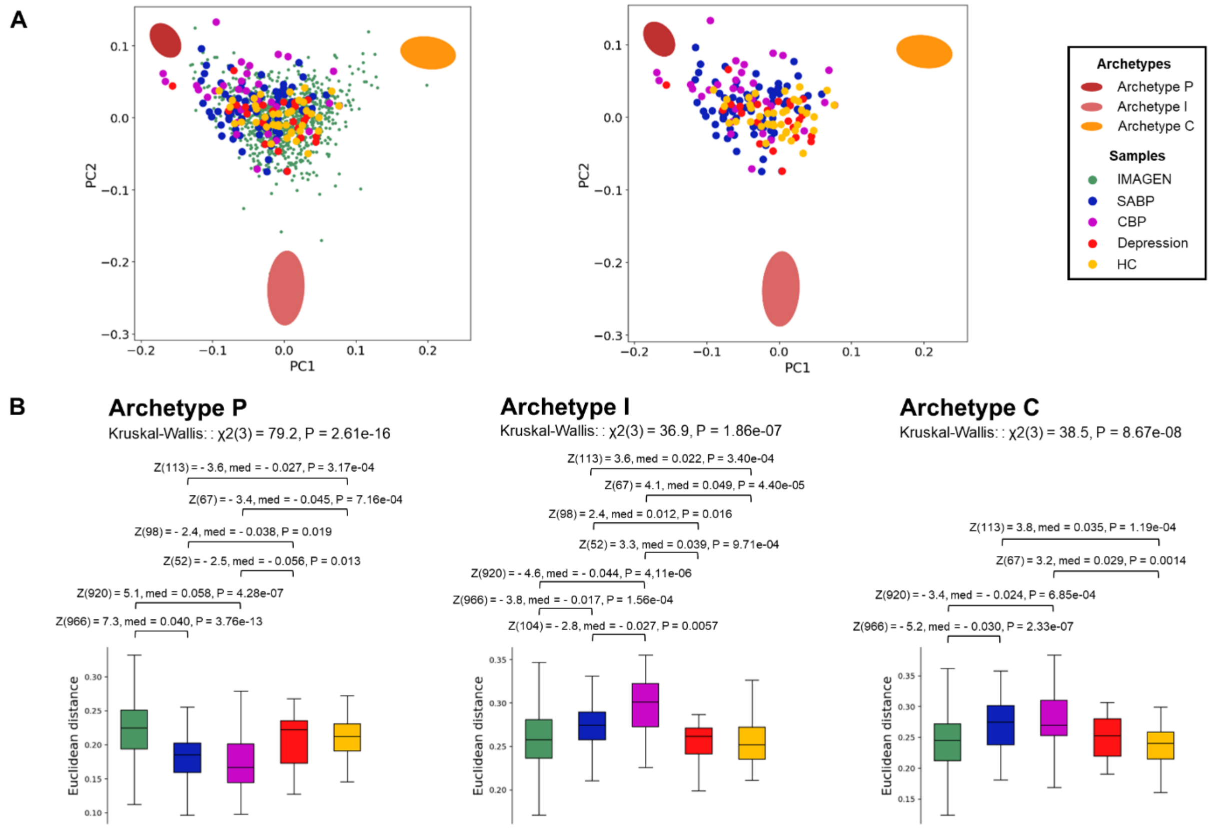


**Fig. S1 | Comparison between the sample from the general population, the three clinical samples (SABP, CBP, Depression) and a sample of healthy controls acquired alongside the clinical pain samples. (A)** All the five samples are shown along with the three archetypes in the 2D space represented by the first two principal components (PCs) of the adjusted proportion of variance of the 19 RSNs. On the left, the IMAGEN sample is included, while on the right it is excluded for a better visualization of the other samples. Clinical pain samples are closer to Archetype P and further from Archetypes I and C with respect to both IMAGEN and the group of healthy controls. **(B)** For each archetype, differences in the Euclidean distances across samples are tested through Kruskal-Wallis tests, followed by post-hoc Mann-Whitney U-tests (FDR < 0.05). SABP = subacute back pain; CBP = chronic back pain; HC = healthy controls; Z = z-score; P = p-value; $\chi2$ = chi-square; PC = principal component.

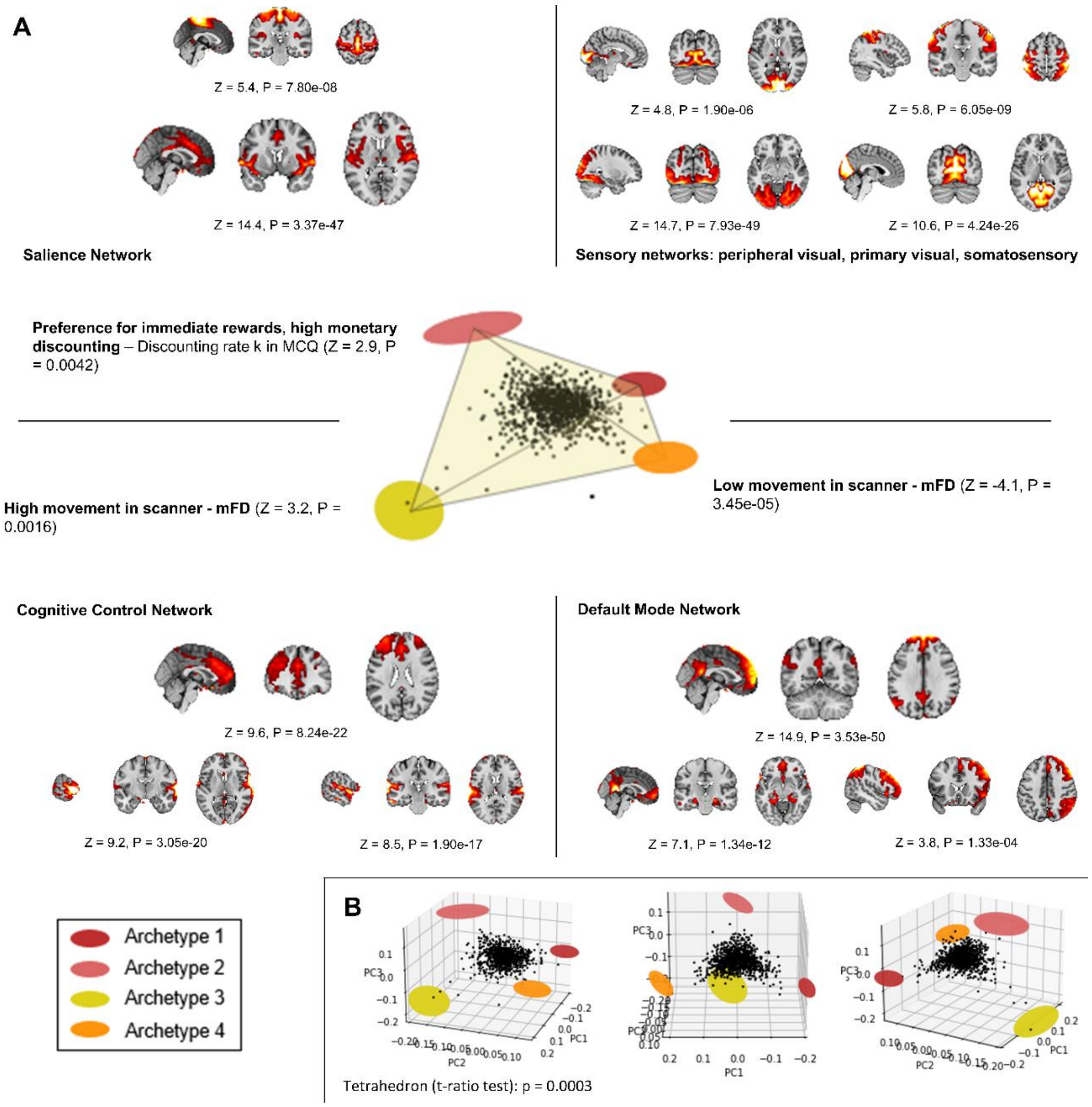


**Fig. S2 | Visualization of the four-archetype solution. (A)** Key findings from the enrichment analysis involving both RSNs and cognitive-behavioral features are displayed for each archetype. Archetype 1 corresponds to Archetype P, Archetype 4 to Archetype C, and Archetype I splits into Archetypes 2 and 3. Archetype 2 is mainly characterized by the salience network and preference for immediate rewards, while Archetype 3 is mainly characterized by a cognitive control network and high in-scanner movement. Consistent with our primary analyses, this suggests that Archetype I optimizes specific aspects of impulsivity. Learning from punishments, represented by higher scores in the Passive Avoidance Learning Paradigm, was still highest near Archetype P, but did not pass the FDR correction (Z(890) = 2.3, P = 0.021). Archetype C was associated with low in-scanner movement, which is in line with our primary analyses where Archetype C was optimized for selfcontrol. In this case, conscientiousness was still highest near Archetype C, but did not pass the FDR correction (Z(890) = 2.2, P = 0.031). Monetary discounting was measured by the discounting rate *k* in the Monetary Choice Questionnaire. For detailed results on all features, refer to Data S1. **(B)** The entire IMAGEN sample (*n* = 892) is plotted within the space defined by the first three PCs of the adjusted proportion of variance of the RSNs. The data are significantly confined in a tetrahedron, as shown by the t-ratio test. The centers of the colored ellipsoids indicate the expected positions of the archetypes, while the ellipsoid shapes represent the associated error. Z = z-score; P = p-value; PC = principal component; mFD = mean framewise displacement; MCQ = Monetary Choice Questionnaire.

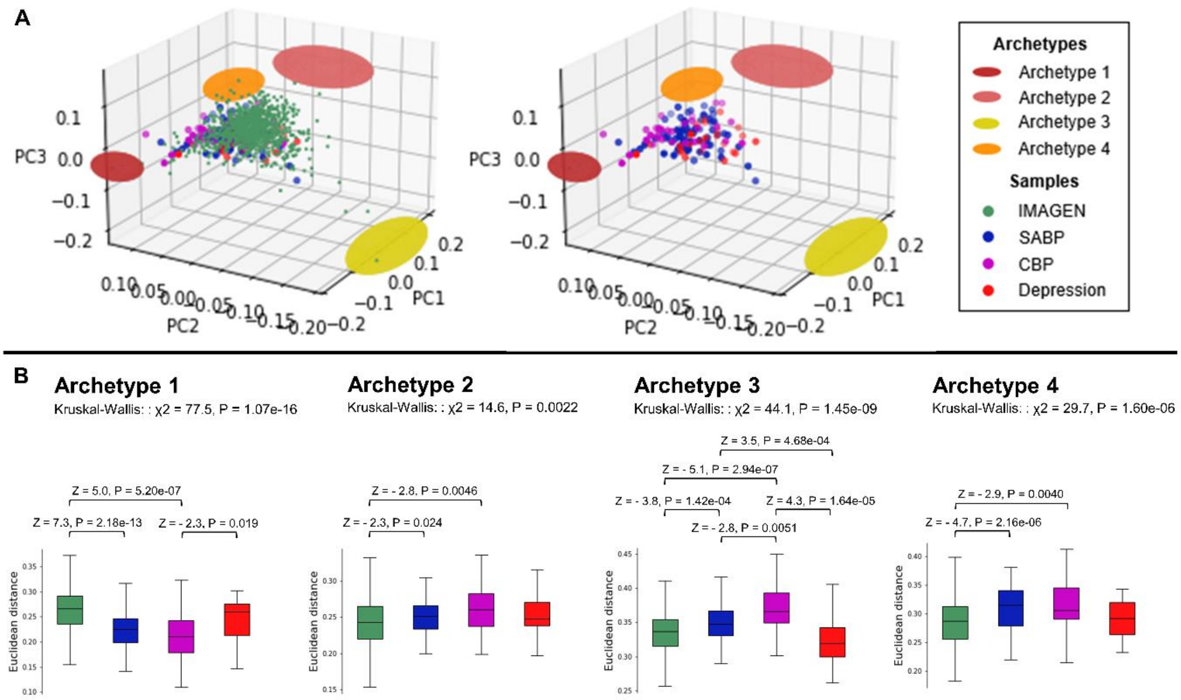


**Fig. S3 | Comparison between the sample from the general population and the three clinical samples (SABP, CBP, Depression) in the solution with four archetypes. (A)** All the four samples are shown along with the four archetypes in the 3D space represented by the first three principal components (PCs) of the adjusted proportion of variance of the 19 RSNs. On the left, the IMAGEN sample is included, while on the right it is excluded for a better visualization of the clinical samples. Clinical pain samples are closer to Archetype P (here called Archetype 1) with respect to IMAGEN. **(B)** For each archetype, differences in the Euclidean distances across samples are tested through Kruskal-Wallis tests, followed by post-hoc Mann-Whitney U-tests (FDR < 0.05). SABP = subacute back pain; CBP = chronic back pain; Z = z-score; P = p-value; χ2 = chi-square; PC = principal component.

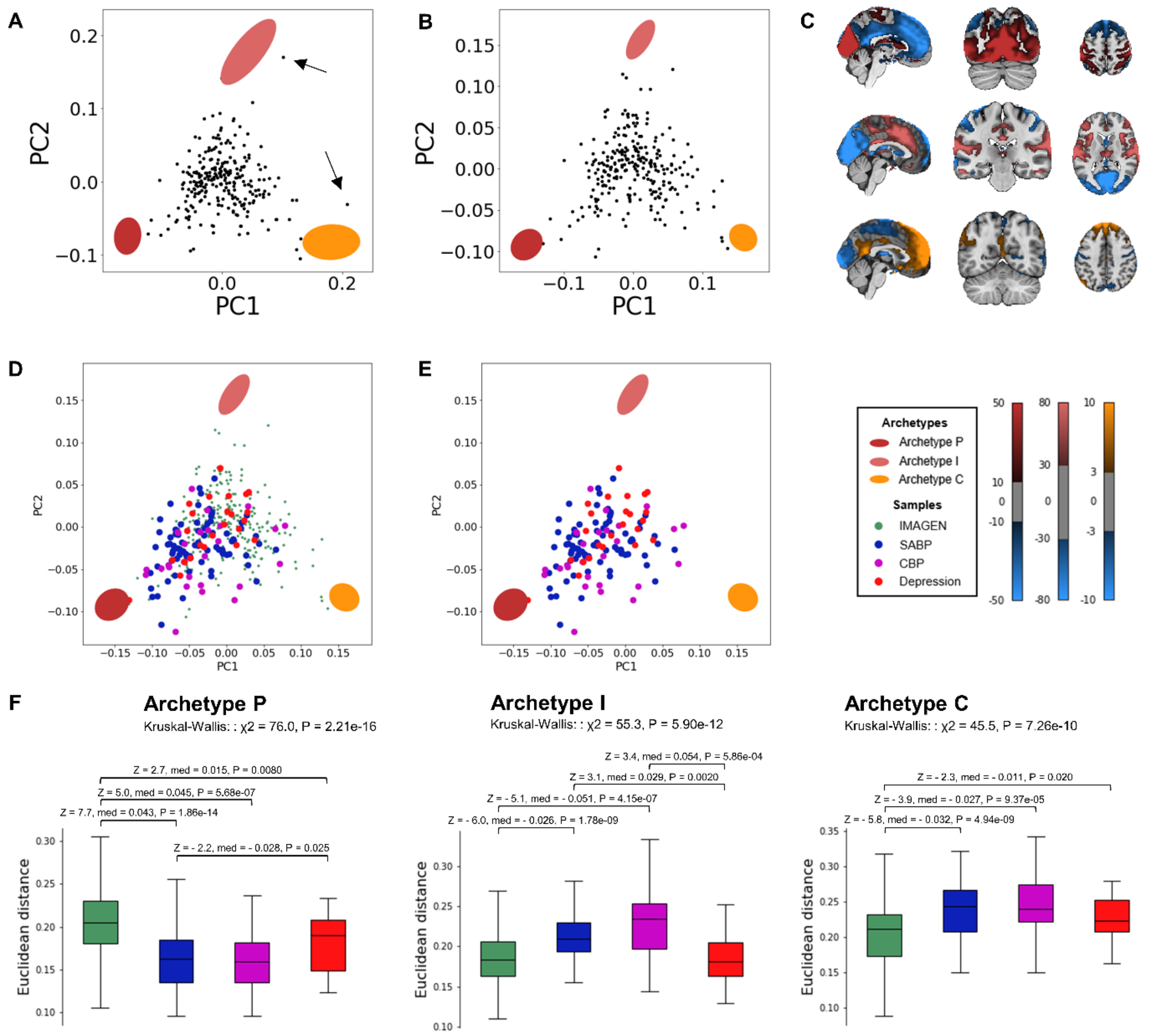


**Fig. S4 | Control analyses using only the subjects from IMAGEN without a field map (*n* = 260).** **(A)** Two outliers (arrows) increased the uncertainty on the positions of Archetypes I and C influencing the t-ratio test, which was not significant (P = 0.13). **(B)** Excluding these two outliers resulted in a significant triangle (P = 0.0068). **(C)** The brain maps of the archetypes are similar to those presented in the main Results, with Archetype P associated with the visual networks, Archetype I with the ACC-Ins salience network, and Archetype C with the default mode network. The PALP score during the PP condition does not pass the FDR correction (FDR<0.1), but it is still maximal close to Archetype P (Z(256) = 2.5, med = 30, P = 0.014). The clinical samples were projected onto the space where the archetypes were defined, with **(D)** and without **(E)** the IMAGEN sample. **(F)** Distances from the archetypes were compared across samples (FDR < 0.05). All the clinical samples (SABP, CBP, and Depression) mapped closer to Archetype P than the IMAGEN sample. SABP = subacute back pain;

CBP = chronic back pain; Z = z-score; med = median difference; P = p-value; $\chi 2$ = chi-square; PC = principal component; PALP = passive avoidance learning paradigm; PP = punishment-punishment.

### Supplementary Tables

**Table S1 | The behavioral constructs used in the enrichment analysis of the IMAGEN sample.** The table shows the constructs used, the questionnaires and tests they are taken from, the percentage of missing values, and the respective references.

| Construct | Questionnaire/Test | Missing values (%) | References |
|---|---|---|---|
| Non-painful somatic symptoms | CSI – Sum of items 2, 4, 7, 8, 9, 10, 11, 12, 13, 14, 15, and from 17 to 32 | 23.0 | Meesters et al., 2003 *(74)* |
| Max pain intensity | CSI – Max score across pain-related items (1, 3, 5, 6, 16, 33, 34, 35) | 23.0 | Meesters et al., 2003 *(74)* |
| Number of painful sites | CSI – Number of items with a score greater than 0 among the pain-related items (1, 3, 5, 6, 16, 33, 34, 35) | 23.0 | Meesters et al., 2003 *(74)* |
| Depressive symptoms | ADRS | 5.0 | Revah-Levy et al., 2007 *(75)* |
| Anxiety symptoms | ANXDX | 11.8 | Wittchen and Pfister, 1997 *(76)* |
| Neuroticism, Extraversion, Openness, Agreeableness, Conscientiousness | NEO | 2.0 | Costa and McCrae, 1992 *(77)* |
| Rumination | RRS | 23.2 | Treynor et al., 2003 *(78)* |
| Emotional abuse, Physical abuse, Sexual abuse, Emotional neglect, Physical neglect | CTQ-SF | 7.2 | Bernstein et al., 2003 *(79)* |
| Average school grades | ESPAD | 2.6 | Hibell et al., 1997 *(80)* |
| Attention impulsivity, Motor impulsivity, Planning impulsivity | BIS-11 | 12.1 | Patton et al., 1995 *(81)* |
| Delay aversion, Deliberation time, Overall proportion bet, Quality of decision making, Risk adjustment, Risk taking | CGT | 1.5 | CANTAB®, Cambridge Cognition, 2019 |

| | | | |
|---|---|---|---|
| Monetary discounting (discounting rate k) | MCQ | 2.9 | Kirby et al., 1999 *(82)* |
| Score PP, Score RP, Score RR, Omission errors PP, Omission errors RP, Omission errors RR, Commission errors PP, Commission errors RP, Commission errors RR, Reaction times PP, Reaction times RP, Reaction times RR | PALP | 4.5, 4.5, 5.0, 4.5, 4.5, 5.0, 4.5, 4.5, 5.0, 4.5, 4.5, 5.2 | Arnett and Newman, 2000 *(83)* |
| Hopelessness, Anxiety sensitivity, Impulsivity, Sensation seeking | SURPS | 3.0 | Woicik et al., 2009 *(84)* |

Abbreviations: ADRS = Adolescent Depression Rating Scale; ANXDX = Anxiety Screening, out of the Composite Diagnostic Interview DIA-X/M-CIDI; BIS-11 = Barratt Impulsiveness Scale; CGT = Cambridge Gambling Task; CSI = Children's Somatization Inventory; CTQ-SF = Childhood Trauma Questionnaire; ESPAD = European School Survey Project on Alcohol and Drugs; MCQ = Monetary Choice Questionnaire; NEO = Neuroticism-Extraversion-Openness Personality Inventory; PALP = Passive Avoidance Learning Paradigm; RRS = Ruminative Responses Scale; SURPS = Substance Use Risk Profile Scale; PP = punishment-punishment condition; RP = reward-punishment condition; RR = reward-reward condition.

**Table S2 | Distances from the archetypes are not associated with current pharmacological treatments.** Mann-Whitney U-tests were performed on the 19D Euclidean distances from the archetypes of people treated vs. non-treated with a certain drug, using the SABP sample (*n* = 76). Tests were done only for those medicaments or categories of drugs taken by at least 5 individuals.

| Treatment | Archetype | P | Z | Med |
|---|---|---|---|---|
| **NSAIDs (34/76; Ibuprofen, Aspirin, Diclofenac)** | Archetype P | 0.55 | 0.60 | 0.0044 |
| | Archetype I | 0.26 | 1.12 | 0.012 |
| | Archetype C | 0.28 | -1.08 | -0.016 |
| **Ibuprofen (30/76)** | Archetype P | 0.66 | 0.44 | 0.0028 |
| | Archetype I | 0.22 | 1.24 | 0.012 |
| | Archetype C | 0.20 | -1.27 | -0.022 |
| **Hormonal contraceptives (10/76; birth pills + one person with a progestin-releasing implant)** | Archetype P | 0.22 | 1.22 | 0.012 |
| | Archetype I | 0.88 | -0.15 | 0.0002 |
| | Archetype C | 0.91 | -0.12 | 0.0029 |
| **Aspirin (5/76)** | Archetype P | 0.59 | -0.54 | -0.018 |
| | Archetype I | 0.25 | 1.15 | 0.010 |
| | Archetype C | 0.36 | 0.92 | 0.015 |
| **Paracetamol (5/76)** | Archetype P | 0.90 | 0.13 | 0.0026 |
| | Archetype I | 0.53 | -0.63 | -0.011 |
| | Archetype C | 0.52 | -0.65 | -0.013 |
| **L-Thyroxin (5/76)** | Archetype P | 0.62 | -0.50 | -0.027 |
| | Archetype I | 0.83 | 0.21 | 0.0063 |
| | Archetype C | 0.74 | 0.34 | 0.0071 |

Abbreviations: P = p-value; Z = z-value; Med = median difference; NSAID = nonsteroidal anti-inflammatory drug.

**Table S3 | The behavioral constructs used in the enrichment analysis of the clinical pain samples.** The table shows the constructs used, the questionnaires and tests they are taken from, the percentage of missing values, and the respective references.

| Construct | Questionnaire/Test | Missing values (%) | References |
|---|---|---|---|
| Pain severity, Interference, Affective distress, Support, Life control, Punishing responses (by significant other), Solicitous responses (by significant other), Distracting responses (by significant other), Activities away from home and social activities, Household chores, Outdoor work | German version of the MPI | 9.0 | Flor et al., 1990 *(85)*; Kerns, Turk, and Rudy, 1985 *(86)* |
| Helplessness, Resourcefulness | PRCS | 26.0 | Flor, Behle, and Birbaumer, 1993 *(87)* |
| Active coping, Catastrophizing | PRSS | 25.4 | Flor, Behle, and Birbaumer, 1993 *(87)* |
| Depressive symptoms, Anxiety symptoms | HADS | 9.0 | Zigmond and Snaith, 1983 *(88)* |

Abbreviations: MPI = West Haven-Yale Multidimensional Pain Inventory; PRCS = Pain-Related Control Scale; PRSS = Pain-Related Self-Statements Scale; HADS = Hospital Anxiety and Depression Scale.

**Table S4 | Demographics for each center of the IMAGEN sample used in the analyses (*n* = 892).**

| | **Berlin** | **Dresden** | **Dublin** | **Hamburg** | **London** | **Mannheim** | **Nottingham** | **Paris** |
|---|---|---|---|---|---|---|---|---|
| **Sex (F)** | 0.61 | 0.51 | 0.48 | 0.50 | 0.56 | 0.55 | 0.57 | 0.59 |
| **Age** | 19.1 +/- 0.9 | 18.6 +/- 0.5 | 19.3 +/- 0.6 | 18.8 +/- 0.5 | 19.0 +/- 0.6 | 19.0 +/- 0.8 | 18.9 +/- 0.7 | 19.7 +/- 0.7 |

**Table S5 | DAWBA bands of probabilities for diagnoses of mental disorders in the IMAGEN sample.** Probabilities for diagnoses of mental disorders, based on bands defined in the Development and Well-Being Assessment (DAWBA; Goodman et al., 2000) *(89)*. For each disorder, the proportion of subjects in each band is reported. Data are based on the sample used in the analyses.

| P(D) | MDD | GAD | SP | SAD | PD | AG | PTSD | OCD | ED |
|---|---|---|---|---|---|---|---|---|---|
| **< 0.1%** | 0.564 | 0 | 0.656 | 0.745 | 0.967 | 0.910 | 0.887 | 0.832 | 0.628 |
| **≈ 0.5%** | 0.316 | 0.728 | 0.328 | 0.170 | 0.021 | 0.086 | 0.093 | 0.140 | 0.276 |
| **≈ 3%** | 0 | 0.175 | 0 | 0.043 | 0 | 0 | 0.009 | 0.015 | 0.084 |
| **≈ 15%** | 0.086 | 0.079 | 0.009 | 0.029 | 0.006 | 0.005 | 0.008 | 0.011 | 0 |
| **≈ 50%** | 0.021 | 0.018 | 0.007 | 0.013 | 0.007 | 0 | 0.003 | 0.002 | 0.013 |
| **> 70%** | 0.013 | 0 | 0 | 0 | 0 | 0 | 0 | 0 | 0 |

Abbreviations: P(D) = probability of diagnosis; MDD = major depression; GAD = generalized anxiety disorder; SP = specific phobia; SAD = social anxiety disorder (social phobia); PD = panic disorder; AG = agoraphobia; PTSD = post-traumatic stress disorder; OCD = obsessive compulsive disorder; ED = eating disorders.

**Table S6 | Scanning parameters of the resting-state protocol for each center, from the follow-up 2 of the IMAGEN dataset.**

| | Berlin | Dresden | Mannheim | Hamburg | London | Nottingham | Dublin | Paris |
|---|---|---|---|---|---|---|---|---|
| **Scanner** | SIEMENS MAGNETOM Verio Syngo MR B17 | SIEMENS MAGNETOM TrioTim Syngo MR B17 | SIEMENS MAGNETOM TrioTim Syngo MR B17 | SIEMENS MAGNETOM TrioTim Syngo MR B17 | General Electric | PHILIPS | PHILIPS | SIEMENS MAGNETOM TrioTim Syngo MR B17 |
| **TR (s)** | 2.2 | 2.2 | 2.2 | 2.2 | 2.2 | 2.2 | 2.2 | 2.2 |
| **TE (ms)** | 30 | 30 | 30 | 30 | 30 | 30 | 30 | 30 |
| **Matrix size** | 64 x 64 | 64 x 64 | 64 x 64 | 64 x 64 | 64 x 64 | 64 x 64 | 64 x 64 | 64 x 64 |
| **Acceleration factor** | 2 | 2 | 2 | 2 | 2 | 2 | 2 | 2 |
| **No. of slices** | 40 | 40 | 40 | 40 | 40 | 40 | 40 | 40 |
| **Slice thickness (mm)** | 2.4 | 2.4 | 2.4 | 2.4 | 2.4 | 2.4 | 2.4 | 2.4 |
| **Slice gap (mm)** | 1.0 | 1.0 | 1.0 | 1.0 | 1.0 | 1.0 | 1.0 | 1.0 |
| **Slice encoding direction** | Desc. | Desc. | Desc. | Desc. | Desc. | Desc. | Desc. | Desc. |
| **Slice acquisition mode** | Seq. | Seq. | Seq. | Seq. | Seq. | Seq. | Seq. | Seq. |
| **Phase encoding direction** | A>>P | P>>A | P>>A | P>>A | P>>A | A>>P | P>>A | P>>A |
| **Echo spacing (ms)** | 0.52 | 0.58 | 0.58 | 0.58 | 0.424 | 0.5842 | 0.5842 | 0.58 |
| **FoV ($mm^2$)** | 220 x 220 | 220 x 220 | 220 x 220 | 220 x 220 | 220 x 220 | 220 x 220 | 220 x 220 | 220 x 220 |
| **Flip angle (degrees)** | 75 | 75 | 75 | 75 | 75 | 75 | 75 | 75 |
| **Field map echo time difference (ms)** | 2.46 | 2.46 | 2.46 | 2.46 | 2.27 | 5 | 5 | 2.46 |

Abbreviations: TR = time of repetition; TE = time of echo; Desc. = descending; Seq. = sequential; A = anterior; P = posterior; FoV = field of view.

**Table S7 | Scanning parameters of the MPRAGE structural MRI for each center, from the follow-up 2 of the IMAGEN dataset.**

| | Berlin | Dresden | Mannheim | Hamburg | London | Nottingham | Dublin | Paris |
|---|---|---|---|---|---|---|---|---|
| **Scanner** | SIEMENS MAGNETOM Verio Syngo MR B17 | SIEMENS MAGNETOM TrioTim Syngo MR B17 | SIEMENS MAGNETOM TrioTim Syngo MR B17 | SIEMENS MAGNETOM TrioTim Syngo MR B17 | General Electric | PHILIPS | PHILIPS | SIEMENS MAGNETOM TrioTim Syngo MR B17 |
| **TR (s)** | 2.3 | 2.3 | 2.3 | 2.3 | 2.3 | 2.3 | 2.3 | 2.3 |
| **TE (ms)** | 2.8 | 2.8 | 2.8 | 2.8 | 2.8 | 2.8 | 2.8 | 2.8 |
| **Matrix size** | 256 x 256 | 256 x 256 | 256 x 256 | 256 x 256 | 256 x 256 | 256 x 256 | 256 x 256 | 256 x 256 |
| **No. of slices** | 160 | 160 | 160 | 160 | 170 | 137 | 137 | 160 |
| **Slice thickness (mm)** | 1.1 | 1.1 | 1.1 | 1.1 | 1.1 | 1.1 | 1.1 | 1.1 |
| **FoV ($mm^2$)** | 263 x 280 | 263 x 280 | 263 x 280 | 263 x 280 | 263 x 280 | 263 x 281 | 263 x 281 | 263 x 280 |
| **Flip angle (degrees)** | 9 | 9 | 9 | 9 | 8 | 9 | 9 | 9 |

Abbreviations: TR = time of repetition; TE = time of echo; FoV = field of view.

### Supplementary Methods

#### Preprocessing of fMRI data

The boilerplate from fMRIPrep, detailing the preprocessing steps, is provided below.

<u>fMRIPrep boilerplate:</u>

Results included in this manuscript come from preprocessing performed using *fMRIPrep* 21.0.1 (@fmriprep1; @fmriprep2; RRID:SCR_016216), which is based on *Nipype* 1.6.1 (@nipype1; @nipype2; RRID:SCR_002502).

Preprocessing of B0 inhomogeneity mappings:

A total of 1 fieldmaps were found available within the input BIDS structure for this particular subject. A *B0* nonuniformity map (or *fieldmap*) was estimated from the phase-drift map(s) measure with two consecutive GRE (gradient-recalled echo) acquisitions. The corresponding phase-map(s) were phase-unwrapped with `prelude` (FSL 6.0.5.1:57b01774).

Anatomical data preprocessing:

A total of 1 T1-weighted (T1w) images were found within the input BIDS dataset. The T1-weighted (T1w) image was corrected for intensity non-uniformity (INU) with `N4BiasFieldCorrection` [@n4], distributed with ANTs 2.3.3 [@ants, RRID:SCR_004757], and used as T1w-reference throughout the workflow. The T1w-reference was then skull-stripped with a *Nipype* implementation of the `antsBrainExtraction.sh` workflow (from ANTs), using OASIS30ANTs as target template. Brain tissue segmentation of cerebrospinal fluid (CSF), white-matter (WM) and gray-matter (GM) was performed on the brain-extracted T1w using `fast` [FSL 6.0.5.1:57b01774, RRID:SCR_002823, @fsl_fast]. Volume-based spatial normalization to two standard spaces (MNI152NLin2009cAsym, MNI152NLin6Asym) was performed through nonlinear registration with `antsRegistration` (ANTs 2.3.3), using brain-extracted versions of both T1w reference and the T1w template. The following templates were selected for spatial normalization: *ICBM 152 Nonlinear Asymmetrical template version 2009c* [@mni152nlin2009casym, RRID:SCR_008796; TemplateFlow ID: MNI152NLin2009cAsym], *FSL's MNI ICBM 152 non-linear 6th Generation Asymmetric Average Brain Stereotaxic Registration Model* [@mni152nlin6asym, RRID:SCR_002823; TemplateFlow ID: MNI152NLin6Asym].

Functional data preprocessing:

First, a reference volume and its skull-stripped version were generated using a custom methodology of *fMRIPrep*. Head-motion parameters with respect to the BOLD reference (transformation matrices, and six corresponding rotation and translation parameters) are estimated before any spatiotemporal filtering using `mcflirt` [FSL 6.0.5.1:57b01774, @mcflirt]. The estimated *fieldmap* was then aligned with rigid-registration to the target EPI (echo-planar imaging) reference run. The field coefficients were mapped on to the reference EPI using the transform. BOLD runs were slice-time corrected to 1.07s (0.5 of slice acquisition range 0s-2.15s) using `3dTshift` from AFNI  [@afni, RRID:SCR_005927]. The

BOLD reference was then co-registered to the T1w reference using `mri_coreg` (FreeSurfer) followed by `flirt` [FSL 6.0.5.1:57b01774, @flirt] with the boundary-based registration [@bbr] cost-function. Co-registration was configured with six degrees of freedom. Several confounding time-series were calculated based on the *preprocessed BOLD*: framewise displacement (FD), DVARS and three region-wise global signals. FD was computed using two formulations following Power (absolute sum of relative motions, @power_fd_dvars) and Jenkinson (relative root mean square displacement between affines, @mcflirt). FD and DVARS are calculated for each functional run, both using their implementations in *Nipype* [following the definitions by @power_fd_dvars]. The three global signals are extracted within the CSF, the WM, and the whole-brain masks. Additionally, a set of physiological regressors were extracted to allow for component-based noise correction [*CompCor*, @compcor]. Principal components are estimated after high-pass filtering the *preprocessed BOLD* time-series (using a discrete cosine filter with 128s cut-off) for the two *CompCor* variants: temporal (tCompCor) and anatomical (aCompCor). tCompCor components are then calculated from the top 2% variable voxels within the brain mask. For aCompCor, three probabilistic masks (CSF, WM and combined CSF+WM) are generated in anatomical space. The implementation differs from that of Behzadi et al. in that instead of eroding the masks by 2 pixels on BOLD space, the aCompCor masks are subtracted a mask of pixels that likely contain a volume fraction of GM. This mask is obtained by thresholding the corresponding partial volume map at 0.05, and it ensures components are not extracted from voxels containing a minimal fraction of GM. Finally, these masks are resampled into BOLD space and binarized by thresholding at 0.99 (as in the original implementation). Components are also calculated separately within the WM and CSF masks. For each CompCor decomposition, the *k* components with the largest singular values are retained, such that the retained components' time series are sufficient to explain 50 percent of variance across the nuisance mask (CSF, WM, combined, or temporal). The remaining components are dropped from consideration. The head-motion estimates calculated in the correction step were also placed within the corresponding confounds file. The confound time series derived from head motion estimates and global signals were expanded with the inclusion of temporal derivatives and quadratic terms for each [@confounds_satterthwaite_2013]. Frames that exceeded a threshold of 0.5 mm FD or 1.5 standardized DVARS were annotated as motion outliers. The BOLD time-series were resampled into standard space, generating a *preprocessed BOLD run in MNI152NLin2009cAsym space*. First, a reference volume and its skull-stripped version were generated using a custom methodology of *fMRIPrep*. Automatic removal of motion artifacts using independent component analysis [ICA-AROMA, @aroma] was performed on the *preprocessed BOLD on MNI space* time-series after removal of non-steady state volumes and spatial smoothing with an isotropic, Gaussian kernel of 6mm FWHM (full-width half-maximum). Corresponding "non-aggressively" denoised runs were produced after such smoothing. Additionally, the "aggressive" noise-regressors were collected and placed in the corresponding confounds file. All resamplings can be performed with *a single interpolation step* by composing all the pertinent transformations (i.e. head-motion transform matrices, susceptibility distortion correction when available, and co-registrations to anatomical and output spaces). Gridded (volumetric) resamplings were performed using `antsApplyTransforms` (ANTs), configured with

Lanczos interpolation to minimize the smoothing effects of other kernels [@lanczos]. Non-gridded (surface) resamplings were performed using `mri_vol2surf` (FreeSurfer).

Many internal operations of *fMRIPrep* use *Nilearn* 0.8.1 [@nilearn, RRID:SCR_001362], mostly within the functional processing workflow. For more details of the pipeline, see [the section corresponding to workflows in *fMRIPrep*'s documentation](https://fmriprep.readthedocs.io/en/latest/workflows.html "FMRIPrep's documentation").
